# Disentangling positive vs. relaxed selection in animal mitochondrial genomes

**DOI:** 10.1101/2022.10.05.510972

**Authors:** Kendra D. Zwonitzer, Erik N. K. Iverson, James J. Sterling, Ryan J. Weaver, Bradley A. Maclaine, Justin C. Havird

**Author notes:** These authors contributed equally to this work (Kendra D. Zwonitzer, Erik N. K. Iverson).

## Abstract

Disentangling different types of selection is a common goal in molecular evolution. Elevated *d*_N_/*d*_S_ ratios (the ratio of nonsynonymous to synonymous substitution rates) in focal lineages are often interpreted as signs of positive selection. Paradoxically, relaxed purifying selection can also result in elevated *d*_N_/*d*_S_ ratios, but tests to distinguish these two causes are seldomly implemented. Here, we reevaluated seven case studies describing elevated *d*_N_/*d*_S_ ratios in animal mtDNA and their accompanying hypotheses regarding selection. They included flightless vs. flighted lineages in birds, bats, and insects, and physiological adaptations in snakes, two groups of electric fishes, and primates. We found that elevated *d*_N_/*d*_S_ ratios were often not caused by the predicted mechanism, and we sometimes found strong support for the opposite mechanism. We discuss reasons why energetic hypotheses may be confounded by other selective forces acting on mtDNA and caution against overinterpreting molecular “spandrels”, including elevated *d*_N_/*d*_S_ ratios.

## Introduction

Detecting different patterns of selection using sequence data is a key goal of molecular evolution. A popular, simple tool in such studies is the *d*_N_/*d*_S_ ratio: the ratio of nonsynonymous to synonymous substitution rates. An abundance of nonsynonymous changes (*d*_N_/*d*_S_ > 1) may indicate positive selection for a gene to undergo adaptive amino acid replacement, while *d*_N_/*d*_S_ ratios near 1 suggest neutral evolution, and *d*_N_/*d*_S_ ratios < 1 indicate purifying selection (Hughes and Nei 1989). Researchers often employ “branch models” to estimate *d*_N_/*d*_S_ ratios across a phylogeny (Yang and Nielsen 1998), with elevated *d*_N_/*d*_S_ ratios in focal lineages suggesting intensified positive selection. Paradoxically, elevated *d*_N_/*d*_S_ ratios can also indicate relaxed purifying selection owing to increased amino acid changes being tolerated more in focal lineages. However, studies rarely explicitly disentangle relaxed vs. positive selection as causes of elevated *d*_N_/*d*_S_ ratios.

Specifically, elevated *d*_N_/*d*_S_ ratios in mitochondrial DNA (mtDNA) are often implicated as evidence for positive selection on energetics. Despite being originally assumed to be a strictly neutral marker (Ballard and Kreitman 1995; Brown et al. 1979; Lynch 1996), mtDNA variation has been convincingly linked to adaptation across different types of environments (Ballard and Melvin 2010; Ballard and Whitlock 2004; Camus et al. 2017; Cheviron and Brumfield 2009; Dobler et al. 2014; Greenway et al. 2020; Hill 2019; James and Ballard 2003), with 26% of nonsynonymous substitutions in animal mtDNA likely fixed by adaptive evolution (James et al. 2016). Given the myriad roles mitochondria play in cellular metabolism, mtDNA variation may be widely important in animal adaptive evolution (Havird et al. 2019b; Hill et al. 2019). Therefore, it is tempting to speculate that signs of positive selection on mtDNA may be found in any animal lineage that occupies an interesting ecological or metabolic niche. Specific hypotheses can be readily tested due to the availability of complete metazoan mtDNA sequences (Fig. 1) and tools for extracting complete mtDNAs from next-generation sequencing datasets (Al-Nakeeb et al. 2017; Allio et al. 2020; Guo et al. 2013; Nachtigall et al. 2021). Examples of recently tested hypotheses include positive selection on mtDNA for low echolocation frequencies in bats (Zhang et al. 2021) and environmental adaptation in gentoo penguins (Noll et al. 2022).

**Figure 1.**
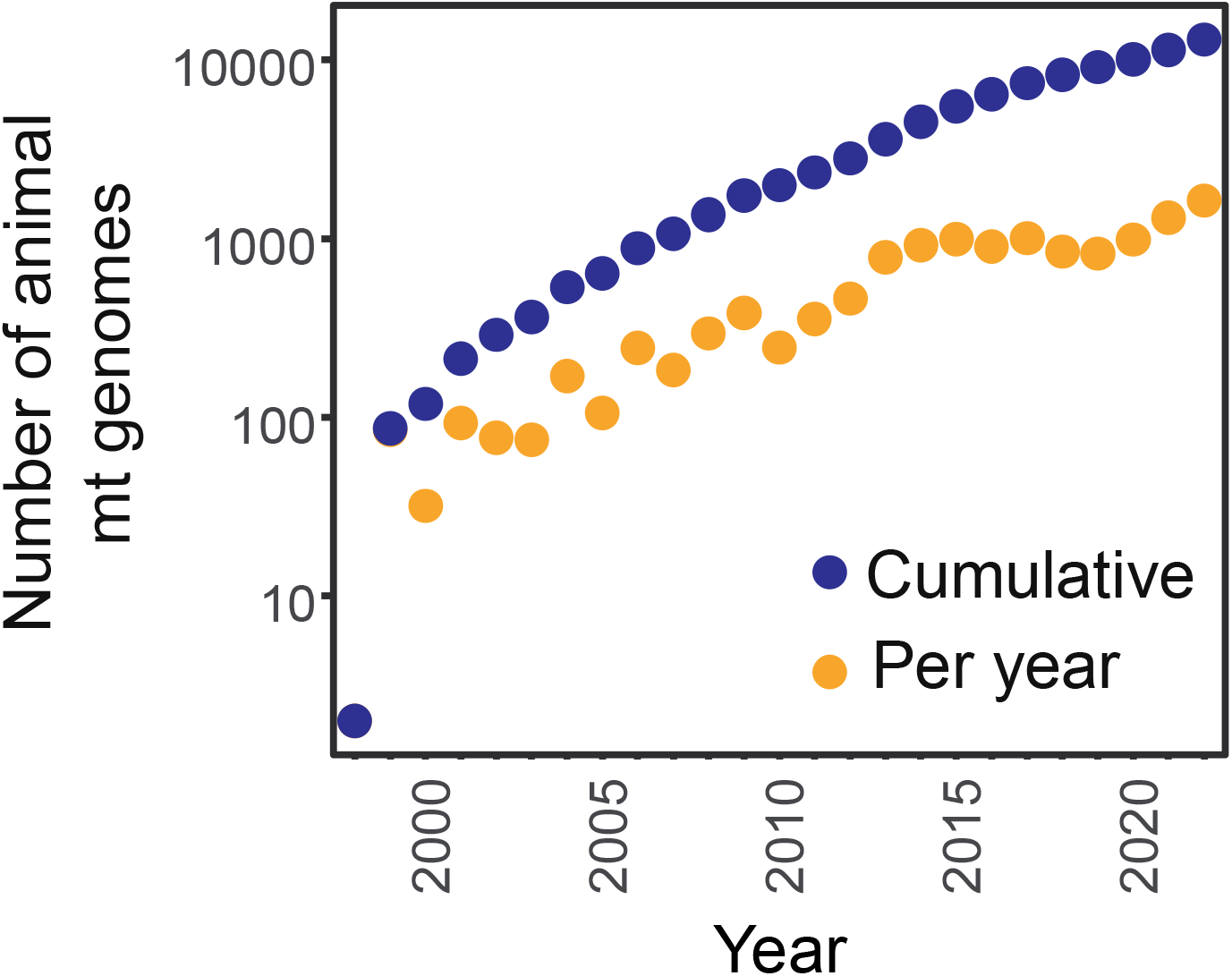
Number of complete animal mtDNA genomes available on NCBI’s organelle database, both per year and cumulative. Accessed on 5 August 2022.

Studies of selection on mtDNA offer a useful example to demonstrate the importance of disentangling positive vs. relaxed selection when interpreting *d*_N_/*d*_S_ results. Relaxed selection on mtDNA (relative to control lineages) is seldom explicitly explored given the core functions of mt genes. However, relatively relaxed selection on mtDNA has been convincingly demonstrated in eusocial shrimps (Chak et al. 2020) and bivalves with doubly-uniparental inheritance of mtDNA (Maeda et al. 2021). These studies used tests that specifically discriminate positive from relaxed selection and found consistent, global signatures of relaxed selection along with elevated *d*_N_/*d*_S_ ratios.

Here, we reanalyzed seven case studies where positive or relaxed selection was hypothesized based on elevated *d*_N_/*d*_S_ ratios from mtDNA datasets. Three cases concerned the evolution/loss of flight and four investigated metabolic innovations. We expanded taxon sampling, calculated branch-specific *d*_N_/*d*_S_ ratios, and used RELAX (Wertheim et al. 2014) to distinguish between positive and relaxed selection on complete mitogenomes. While some conclusions of the original studies may remain valid, our analyses suggest that assuming elevated *d*_N_/*d*_S_ ratios fit a particular narrative without explicitly testing for their causes may lead to erroneous, spandrel-esque conclusions (Gould et al. 1979).

## Materials and Methods

### Positive and relaxed selection in flighted and flightless lineages

The first three case studies we reexamined explored the general hypothesis that mtDNA is under positive selection in flighted compared to flightless lineages because flight is energetically intense. First, mtDNA in bats was hypothesized to be under positive selection compared to flightless mammals (Shen et al. 2010). We therefore compared 77 bats to 58 flightless mammals (including insectivores, carnivores, cetaceans, and ungulates, similar to Shen et al. (2010). For the second and third case studies, mtDNA in flightless birds and insects was hypothesized to be under relatively relaxed selection compared to flighted lineages (Mitterboeck and Adamowicz 2013; Shen et al. 2009). We therefore compared 49 flightless birds to 47 closely related flighted birds and 50 flightless insects to 56 closely-related flighted insects. For birds, species were classified as flighted if they were at least weakly flighted, so tinamous were classified as flighted, while emus and kiwis are considered flightless. Because flight can be more plastic in insects, we categorized ambiguous species as flighted if they produced flighted reproductive females. This is because mitochondria are transmitted only by females and flight can only be selected for via mtDNA changes in females (e.g., Havird et al. 2019a).

### Positive selection for metabolic innovations

The four other case studies hypothesized positive selection on mtDNA in lineages with energetic innovations. First, in snakes compared to lizards, owing to physiological redesign associated with lung volume reduction, consumption of infrequent/large meals, and venom production (Castoe et al. 2008). We therefore compared 21 snakes to 31 closely related lizards. The next two case studies hypothesized positive selection in two independent origins of electric fishes due to their ability to generate electric fields for communication and prey detection (Elbassiouny et al. 2020). Therefore, for South American gymnotiform electric fishes we compared 44 gymnotiforms to 43 closely related characiforms and for the African mormyroid electric fishes we compared 39 mormyroids to 79 closely related osteoglossiforms/clupeiforms. Finally, in the last case study we reexamined the “brain-energy” hypothesis, which states that accelerated evolution of mtDNA in primates reflects adaptive evolution for enhanced brain function (Goldberg et al. 2003; Grossman et al. 2004; Osada and Akashi 2012). We therefore compared 10 primates to 20 closely-related rodents, ungulates, and carnivores. We note that in the bats, snakes, electric fishes, and the primate cases, there is only a single, monophylectic lineage of interest, limiting the phylogenetic power of the analyses compared to the bird and insect cases, where many independent flightless lineages were examined.

### Sequence curation

For each case study we first searched NCBI’s organelle genome database for complete mtDNA sequences from relevant species. In some datasets with limited availability of complete mitogenomes, we used other resources, including the EFISH Genomics 2.0 database for electric fishes (https://efishgenomics.integrativebiology.msu.edu/data/) and Campagna et al. (2019) for flightless birds, where transcriptomes/genomes were searched for mt genes using BLAST (Altschul et al. 1997). For electric fishes and flightless birds, we also *de novo* assembled and annotated complete (or nearly complete) mitogenomes from publicly available Illumina sequencing data. FASTQ files were downloaded from NCBI’s Sequence Read Archive (SRA) and mitogenomes were assembled using MitoFinder v1.4 with default settings (Allio et al. 2020). We therefore report new mitogenomes from six birds, 26 gymnotiforms, and nine mormyroids (GenBank accessions: BK061684-BK061730, Table S1).

### Disentangling positive vs. relaxed selection on mtDNA

We first extracted nucleotide sequences from the 13 mitochondrial protein-coding genes in each mitogenome. For each gene we translated and aligned the resulting amino acid sequences using the proper translation table (either the vertebrate or invertebrate mitochondrial genetic code) for each of the seven datasets outlined above, using Muscle (Edgar 2004) as implemented in MEGA X (Kumar et al. 2018). Resulting alignments were manually corrected by eye and both amino acid and nucleotide sequences were exported as fasta files. Alignments were concatenated to make a “mitogenome” dataset representing all 13 genes. Amino acid sequences from the mitogenome dataset were used to perform phylogenetic reconstruction under maximum likelihood in RAxML v8.2.12 with the gamma WAG model of amino acid substitution and 100 rapid bootstrap replicates (`-f a -# 100 -m PROTGAMMAWAG`) (Stamatakis 2014; Stamatakis et al. 2008). Resulting topologies were rooted according to the original publications referenced above to guide selection analyses. For flightless insects we used a constrained topology at the level of orders based on Misof et al. (2014).

We first used branch-specific models in PAML v4.8 (Yang 2007) to fit two different *d*_N_/*d*_S_ ratios on phylogenetic branches in each dataset (“model = 2”): one for the focal/test lineage(s) (red branches in Figs. 2–3, including terminal and consensus internal branches) and one for reference lineages. Internal branches with descendants in both character states (e.g., flighted and flightless) were coded with the ancestral condition by default (e.g., flighted in insects). We compared the resulting likelihood scores from this model to one where all branches were fit with a single *d*_N_/*d*_S_ ratio (“model = 0”) using a likelihood ratio test. This was performed for each gene individually and the mitogenome dataset in each case study.

**Figure 2.**
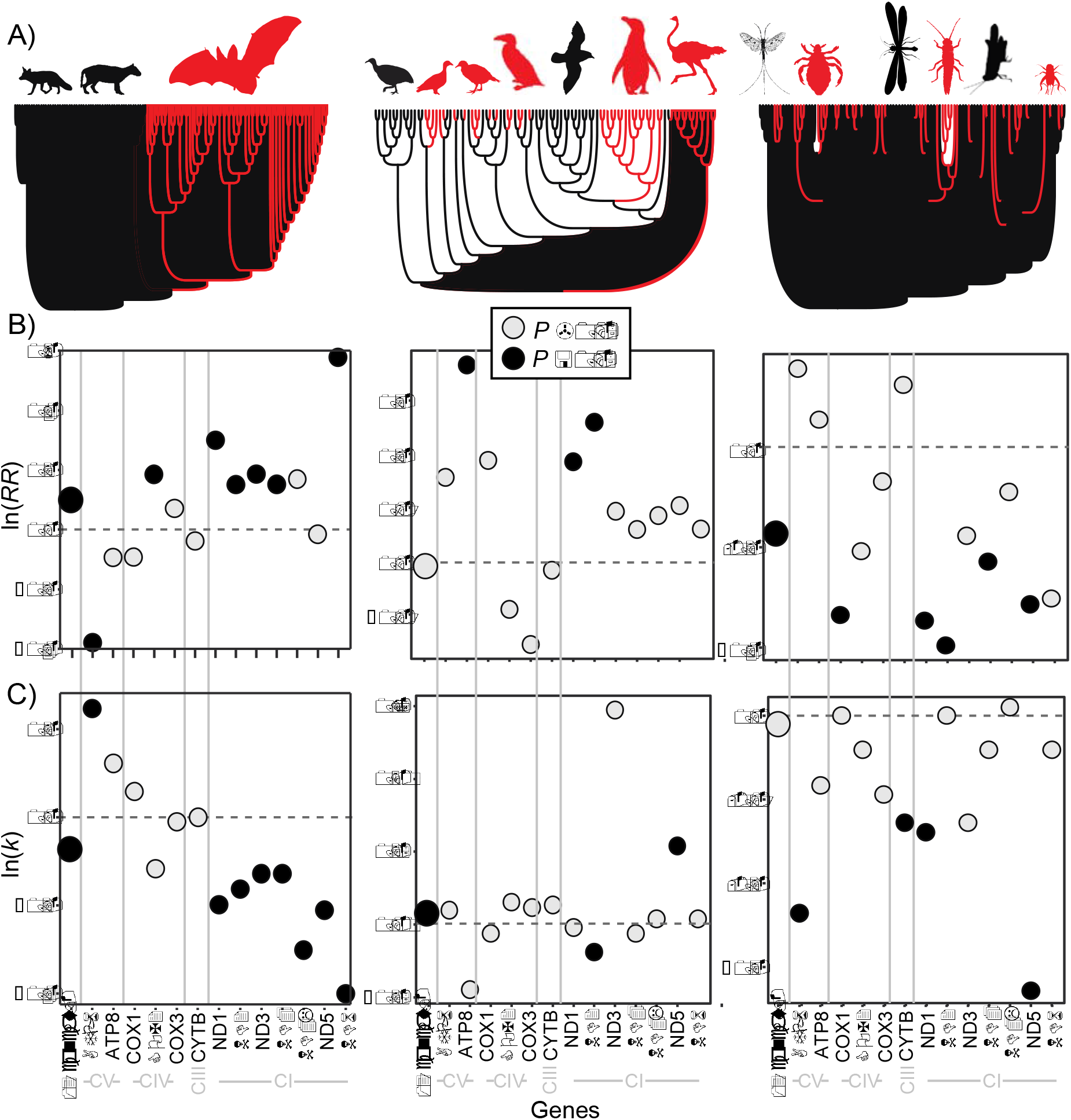
Selection on mitochondrial genes in flightless and flighted lineages. A) Topology showing the number and distribution of lineages (in red; from left to right: bats, flightless birds, and flightless insects) and reference lineages (in black). B) *d*_N_/*d*_S_ ratios in focal lineages compared to reference taxa. Data are presented as the natural log of the response ratio, with numbers above zero indicating higher *d*_N_/*d*_S_ ratios in focal lineages. Statistical significance of a likelihood ratio test compared to a model where only a single *d*_N_/*d*_S_ ratio was used is indicated by black (*P* < 0.05) vs. gray (*P* > 0.05) points. C) Natural log of the *k* parameter from the RELAX analysis where In *k* < 0 indicates relaxed selection in focal lineages and In *k* > 0 indicates positive selection. Statistical significance is indicated as in B). Data are presented for individual mitochondrial genes and the concatenated set of all 13 mitochondrial protein-coding genes.

**Figure 3.**
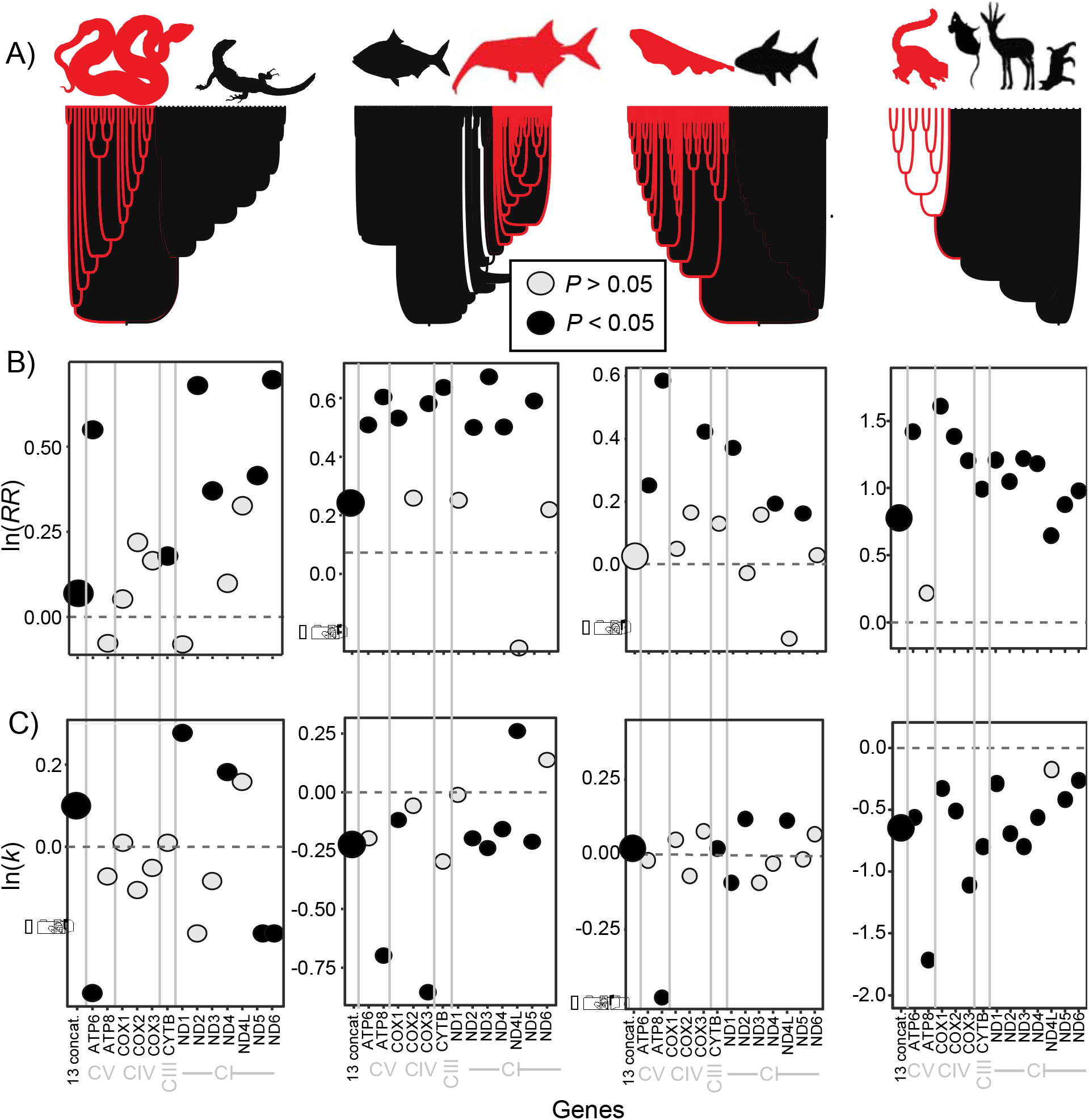
Selection on mitochondrial genes in lineages with energetic innovations. A) Topology showing the number and distribution of focal lineages (in red; from left to right: snakes, mormyroids, gymnotiforms, and primates) and reference lineages (in black). B) *d*_N_/*d*_S_ ratios in focal lineages compared to reference lineages, as in Figure 2. C) RELAX analysis where In *k* < 0 indicates relaxed selection on focal lineages and In *k* > 0 indicates positive selection. Statistical significance and data presentation as in Figure 2.

To distinguish whether positive vs. relaxed selection in focal lineages, we performed analyses using RELAX (Wertheim et al. 2014), both locally within the HyPhy package (Pond et al. 2005) and on the datamonkey webserver (Weaver et al. 2018). Phylogenetic branches were coded as either “test” or “reference” as above (with reference indicating the ancestral condition). Briefly, RELAX compares the distribution of *d*_N_/*d*_S_ ratios across sites in test vs. reference branches, summarized by a *k* parameter, with *k* < 1 suggesting relatively relaxed selection and *k* > 1 suggesting intensified/positive selection in test branches. The statistical significance of the *k* parameter is assessed by comparison to a model where a single distribution of *d*_N_/*d*_S_ ratios is applied across all branches. We performed RELAX analyses on each gene individually and the mitogenome dataset in each case study.

### Data availability

All data analyzed here are publicly available and accession numbers are provided in Table S1. Relevant tree, alignment, and raw data files from this project are available via FigShare (https://figshare.com/articles/dataset/Positive_vs_relaxed_mtDNA_selection/21277536; also happy to upload to Dryad).

## Results

### Inconsistent selection on flighted vs. flightless lineages’ mtDNA

As previously reported, we found increased *d*_N_/*d*_S_ ratios in bats compared to flightless mammals (0.44 vs. 0.40 for the mitogenome, *P* < 0.001, Fig. 2B) (Shen et al. 2010). However, the signal was not consistent: 5 genes showed significantly elevated *d*_N_/*d*_S_ ratios, *ATP6* had a significantly lower *d*_N_/*d*_S_ ratio, and the rest showing non-significant differences. Contrary to the predicted hypothesis, RELAX analyses indicated this pattern was likely due to relaxed, not positive selection on bats (*k* = 0.93 for the mitogenome, *P* < 0.001, Fig. 2C). No individual gene showed the predicted pattern of an elevated *d*_N_/*d*_S_ ratio associated with a signal of positive selection (Fig. 2).

For birds, flightless lineages did not have elevated *d*_N_/*d*_S_ ratios as reported previously (Shen et al. 2009) (0.04 vs. 0.04 for the mitogenome, *P* = 0.717, Fig. 2B). RELAX analyses indicated a weak, but significant trend towards intensified, not relaxed, selection in flightless lineages (*k* = 1.04 for the mitogenome, *P* = 0.034, Fig. 2C). However, only *ND2* showed a significant pattern of elevated *d*_N_/*d*_S_ ratios in flightless taxa (0.06 vs. 0.05, P < 0.001) consistent with relaxed selection (*k* = 0.91, *P* = 0.043).

Contrary to previous results (Mitterboeck and Adamowicz 2013), lower, not elevated, *d*_N_/*d*_S_ ratios were observed in flightless compared to flighted lineages (0.03 vs. 0.04 for the mitogenome, *P* < 0.001, Fig. 2B), which was fairly consistent across genes (Fig. 2B). RELAX analyses were largely statistically non-significant (Fig. 2C), although *k* values in general suggested relaxed selection on flightless lineages. Three individual genes (*ATP6, ATP8*, and *CYTB*) did show the expected patterns of elevated *d*_N_/*d*_S_ ratios associated with signals of relaxed selection.

### Relaxed and positive selection on mtDNA in lineages with energetic innovations

For snakes, we confirmed previously reported elevated *d*_N_/*d*_S_ ratios compared to lizards (0.07 vs. 0.06 for the mitogenome, *P* = 0.002, Fig. 3B) (Castoe et al. 2008), which was consistent across genes (Fig. 3B). As hypothesized, this was likely due to positive selection (*k* = 1.1, *P* <0.001 for the mitogenome, Fig. 3C). However, individual genes were inconsistent. The only two genes with *k* values significantly greater than one (*ND1* and *ND4*) did not have significantly elevated *d*_N_/*d*_S_ ratios. Three individual genes with significantly elevated *d*_N_/*d*_S_ ratios showed *k* values significantly less than one (*ATP6, ND5*, and *ND5*), suggesting relaxed, not positive selection on these genes.

For both gymnotiform and mormyroid electric fishes, there was a clear and consistent trend of elevated *d*_N_/*d*_S_ ratios in electric taxa compared to closely related non-electric fishes, both in the mitogenome (difference in *d*_N_/*d*_S_ ratios of 0.004 – 0.008, *P* < 0.001 for mormyroids, *P* = 0.182 for gymnotiforms, Fig. 3B) and in individual mitochondrial genes (Fig, 3B), as previously reported (Elbassiouny et al. 2020). However, this was not consistent with positive selection via the RELAX results. For mormyroids, there was a clear and consistent pattern of relaxed, not positive, selection (*k* = 0.83, *P* < 0.001 for the mitogenome, similar trends for individual genes, Fig. 3C). For gymnotiforms, there was a less clear pattern, with similar numbers of genes indicating relaxed and positive selection (the mitogenome had a small, but statistically significant effect of positive selection: *k* = 1.02, *P* = 0.024, Fig. 3C).

As suggested previously, primates did show significantly elevated *d*_N_/*d*_S_ ratios across the mitogenome (0.09 vs. 0.04, *P* < 0.001, Fig. 3B) and in nearly all individual genes (up to five-fold for *COX1*). Contradicting the brain-energy hypothesis (Grossman et al. 2004), there was a clear and consistent signal of relaxed, not positive-selection in primates, both in across the mitogenome (*k* = 0.51, P < 0.001, Fig. 3C) and in nearly all individual genes (*ATP8* was the most extreme, being under 5.5 times less intense selection in primates).

## Discussion

Our results suggest that some previous hypotheses may have been based on misinterpreting the underlying causes of elevated *d*_N_/*d*_S_ ratios in mtDNA. We argue that distinguishing between relaxed and intensified positive selection is critical. Of the seven case studies we examined, none showed elevated *d*_N_/*d*_S_ ratios that were consistently attributed to the predicted mechanism, and sometimes the opposite mechanism was convincingly supported. Other times, increased *d*_N_/*d*_S_ ratios in focal lineages were not recovered as in the original analyses, possibly due to increased taxonomic sampling.

Three of the seven case studies we reexamined broadly hypothesize that the evolution of flight drives selection for energetic efficiency because flight is energetically expensive and requires highly efficient mitochondrial function. However, this hypothesis and others related to energetic efficiency in focal lineages, may not be falsifiable based on *d*_N_/*d*_S_ ratios alone. Under this hypothesis, flighted lineages should show higher *d*_N_/*d*_S_ ratios due to positive selection. However, elevated *d*_N_/*d*_S_ ratios in *flightless* taxa due to relaxed selection would also support this hypothesis. Because finding increased *d*_N_/*d*_S_ ratios in either flightless or flighted lineages would support the overall hypothesis, it is difficult to falsify it. Only by distinguishing the cause of increased *d*_N_/*d*_S_ ratios can the hypothesis be properly tested.

While positive selection on mitochondrial genes is an important part of environmental adaptation (Hill 2019), other forces also shape mtDNA evolution. Due to the uniparental inheritance, effective haploidy, and lack of recombination, mtDNA should be under less efficient selection overall than nuclear-encoded genes and have a smaller effective population size (*N_e_*) (Lynch 1996; Lynch and Blanchard 1998; Neiman and Taylor 2009). Differences in *N_e_* may also shape relative selection pressures on mtDNA among lineages (Bazin et al. 2006; Meiklejohn et al. 2007). For example, parasitic lineages often show accelerated mtDNA evolution compared to non-parasitic lineages, possibly owning to reduced *N_e_* (Castro et al. 2002; Jakovlić et al. 2021; Oliveira et al. 2008). Because flightless lineages may have lower dispersal abilities, they may also have lower *N_e_* (Ikeda et al. 2012; McCulloch et al. 2009) and be under relatively relaxed selection, not because they lead a less energetic lifestyle. Many flightless taxa (especially among birds) are also associated with islands and inherently low *N_e_* (Woolfit and Bromham 2005). High *d*_N_/*d*_S_ ratios in mormyroid electric fishes consistent with relaxed selection (Fig. 3) may stem from relatively low *N_e_* compared to reference taxa such as Clupeiformes. Primates also had high *d*_N_/*d*_S_ ratios due to relaxed selection, contrary to the brain-energy hypothesis (Goldberg et al. 2003; Grossman et al. 2004), but consistent with lower *N_e_* in primates compared to reference taxa such as rodents. Other factors independent of energetics also play confounding roles in mtDNA evolution, including longevity (Galtier et al. 2009; Hua et al. 2015; Nabholz et al. 2008), generation time (Thomas et al. 2010), and other cytoplasmic endosymbionts (Hurst and Jiggins 2005).

Multiple factors may shape energetics in complementary or contradictory ways in specific lineages. For example, it has been argued that thermic habit (i.e., endothermy vs. ectothermy) and/or metabolic rate may predict patterns of mtDNA evolution (Rand 1993; Rand 1994). Several of our flightless birds were penguins which may expend more energy on thermoregulation, while others were large, fast-running paleognathes that might have elevated metabolic rates (Fig. 2A). In these taxa, the lower energetic lifestyle stemming from flight loss is confounded with other high-energy adaptations, which may explain why overall *d*_N_/*d*_S_ ratios were similar between flighted and flightless taxa (Fig. 2B). Some studies do attempt to correct for confounding factors. For example, Mitterboeck and Adamowicz (2013) purposely excluded any cases of insect flight loss that were confounded with transitions in life history or population sizes. However, correcting for all confounding factors may be impractical.

Overall, we suggest a cautionary approach when interpreting molecular signatures like *d*_N_/*d*_S_ ratios that are easily calculable, especially as data become increasingly available (Fig. 1). Authors should be skeptical of adaptive “just-so stories” of genetic selection. Unfortunately, tools to explicitly disentangle relaxed vs. positive selection are limited (but see e.g., Crotty et al. 2020). Other standard tools besides *d*_N_/*d*_S_ ratios such as McDonald-Kreitman tests (McDonald and Kreitman 1991) can also have problems in certain contexts (e.g., with mtDNA, Meiklejohn et al. 2007). Selective hypotheses stemming from molecular evolution analyses should ultimately be evaluated through further functional experiments to avoid overinterpretation of molecular “spandrels” (Gould et al. 1979).

## Acknowledgements

We thank the Havird lab for comments on an earlier version of this work. This work was funded by National Institutes of Health 1R35GM142836.

## Notes

https://figshare.com/articles/dataset/Positive_vs_relaxed_mtDNA_selection/21277536

### Competing Interest Statement

The authors have declared no competing interest.

